# Intrinsic fluctuations modulate thalamic gating during the emergence of human conscious perception

**DOI:** 10.64898/2026.10.27.761793

**Authors:** Zepeng Fang, Haifeng Shu, Xin Chen, Anjiao Peng, Yuanyuan Dang, Hulin Zhao, Zheng Li, Jiaxi Zhao, Sixun Yu, Lei Chen, Mingsha Zhang

**Author notes:** ***Co-first authors:*** Zepeng Fang, Haifeng Shu, Xin Chen, Anjiao Peng.

## Abstract

Human conscious perception is not determined solely by external sensory input but is also shaped by moment-to-moment fluctuations in intrinsic brain states. However, how these intrinsic states directly function after stimulus onset to shape the emergence of conscious perception remains unclear. We recorded sEEG activity in thalamus and five other consciousness-related cortical regions, while 32 patients (n= 2,023 recording sites) conscious percept a near-threshold visual stimulus. Preceding the extensively evoked consciousness-related activities, there are early activities (0-200ms after stimulus onset) exclusively appeared in high-order thalamus. These early activities inversely associated with stimulus strength, which reflected an intrinsic state that compensated for the stimulus-evoked response to reach the conscious threshold. These early activities were closer to the critical state in conscious trials, which enables more efficient information processing. These results suggest that intrinsic fluctuations in the high-order thalamus directly modulate thalamic gating during the emergence of conscious perception.

## Introduction

Human conscious perception is not a fixed readout of the external world. Perceptual experiences and underlying neural responses varied from moment-to-moment across repeated presentations of similar sensory input (*1–3*). This phenomenon exposes a fundamental property of brain function, i.e. the brain continuously generates spontaneous fluctuations that reflect its intrinsic state, which shape the upcoming conscious processing of external stimuli.

Probabilistic conscious perception of near-threshold visual stimuli, e.g. sometimes subjectively seen and sometimes not, has provided an important experimental approach for discovering the neural mechanisms underlying consciousness since the 1990s(*4, 5*). Such pheonomenon is proposed to start from the intrinsic fluctuation of brain(*6*). Several EEG studies have shown that prestimulus neural oscillation at alpha-band (8-12 Hz) predict whether a near-threshold stimulus will subsequently be consciously detected(*7–9*). A further intracranial EEG study(*10*) showed that prestimulus alpha activity was negatively correlated with the magnitude of stimulus-evoked high-frequency activity in sensory cortices, suggesting that a hypoexcitable brain state might contribute to the probabilistic process of conscious perception. However, it remains unclear that how intrinsic fluctuations directly link to stimulus-evoked responses and functions immediately after the stimulus onset to initiate brain-wide conscious processing at approximately 200 ms later(*11–14*).

The thalamus is a promising candidate to provide such a mechanism. Beyond relaying sensory signals by first-order thalamic nuclei, higher-order thalamic nuclei play an important role in regulating large-scale brain states (*15*), including neural excitability, interareal communication, and neural coordination. Moreover, human intracranial recordings have recently shown that higher-order thalamus is recruited earlier and more strongly than prefrontal cortex (PFC) during conscious visual perception, and consciousness-related information flows from high-order thalamus to PFC(*16*). These observations suggest that the higher-order thalamus may act as a gate that determines whether weak sensory signals could get access to distributed conscious processing. Yet a critical question remains unresolved: what determines whether the thalamic gate opens? Understanding this mechanism may explain why identical sensory inputs sometimes reach conscious awareness and sometimes do not. Given the functional and anatomical evidence for high-order thalamus, we propose that momentary intrinsic fluctuations in high-order thalamus itself shape the initial opening of the thalamic gate, thereby influencing subsequent stimulus-evoked conscious processing and the emergence of conscious perception.

Most previous studies have relied on scalp electrophysiology or functional magnetic resonance imaging(*2, 6, 9, 17, 18*), which provide limited access to the rapid dynamics of thalamus. Moreover, previous studies usually have generally focused on the prestimulus period due to the limited temporal resolution in the frequency-dependent analysis, and have therefore been unable to elucidate the key mechanisms operating immediately after stimulus onset, when intrinsic and evoked activities directly interact. In addition, distinguishing intrinsic from evoked activity during the poststimulus period is challenging. Thus it remains unclear (i) which types of post-stimulus thalamic activity reflect intrinsic fluctuations (ii) how this activity directly interacts with stimulus-evoked activity, and, most importantly, (iii) which dynamical mechanisms translate these intrinsic fluctuations into probabilistic outcomes of conscious perception.

Criticality is a promising candidate dynamical mechanism for the thalamic gating process during conscious perception. By operating near a critical transition (commonly indexed by branching ratio approaching one), a neural system can achieve an optimal balance between activity propagation and dissipation, enabling both efficient information transmission while keeping a rich repertoire of activity patterns(*19, 20*). As that global conscious states have been associated with such near-critical cortical dynamics(*21–23*), criticality may also play a role in the emergence of conscious perception, i.e. small intrinsic fluctuations in the distance from the near-critical regime could markedly alter the probability that an identical sensory input will rapidly dissipate or propagate and become amplified throughout the brain.

Here we addressed these questions using stereoelectroencephalography (sEEG) while 32 patients performing a visual conscious perception task described by our previous studies(*16, 24, 25*). In this task, conscious perception varied for near-threshold grating stimuli and matched saccadic responses between conscious and unconscious conditions largely minimized the confounding of consciousness-related activity by report-related activity. Combined with the sEEG recordings that sampling at thalamus and other 5 important cortical regions that involved in conscious perception, we were able to thoroughly examine transient intrinsic activity that associated with conscious perception during the peristimulus period with high spatiotemporal resolution.

## Results

### Behavioral results

Subjects’ behavioral performance followed the pattern reported previously, i.e. higher stimulus contrast was associated with more frequent conscious reports, producing a typical sigmoidal curve. In control trials with no grating or high grating contrast, subjects achieved high accuracy (93.78% ± 1.03), indicating that they understood and performed the task well. In trials with near-threshold grating contrast (Fig. 1B, see methods for details), subjects varied in reporting whether they were “conscious” or “unconscious” of the grating. We focused on those near-threshold trials and compared the conscious and unconscious trials with the same contrast within them to isolate consciousness-related changes for further analysis.

**Fig. 1.**
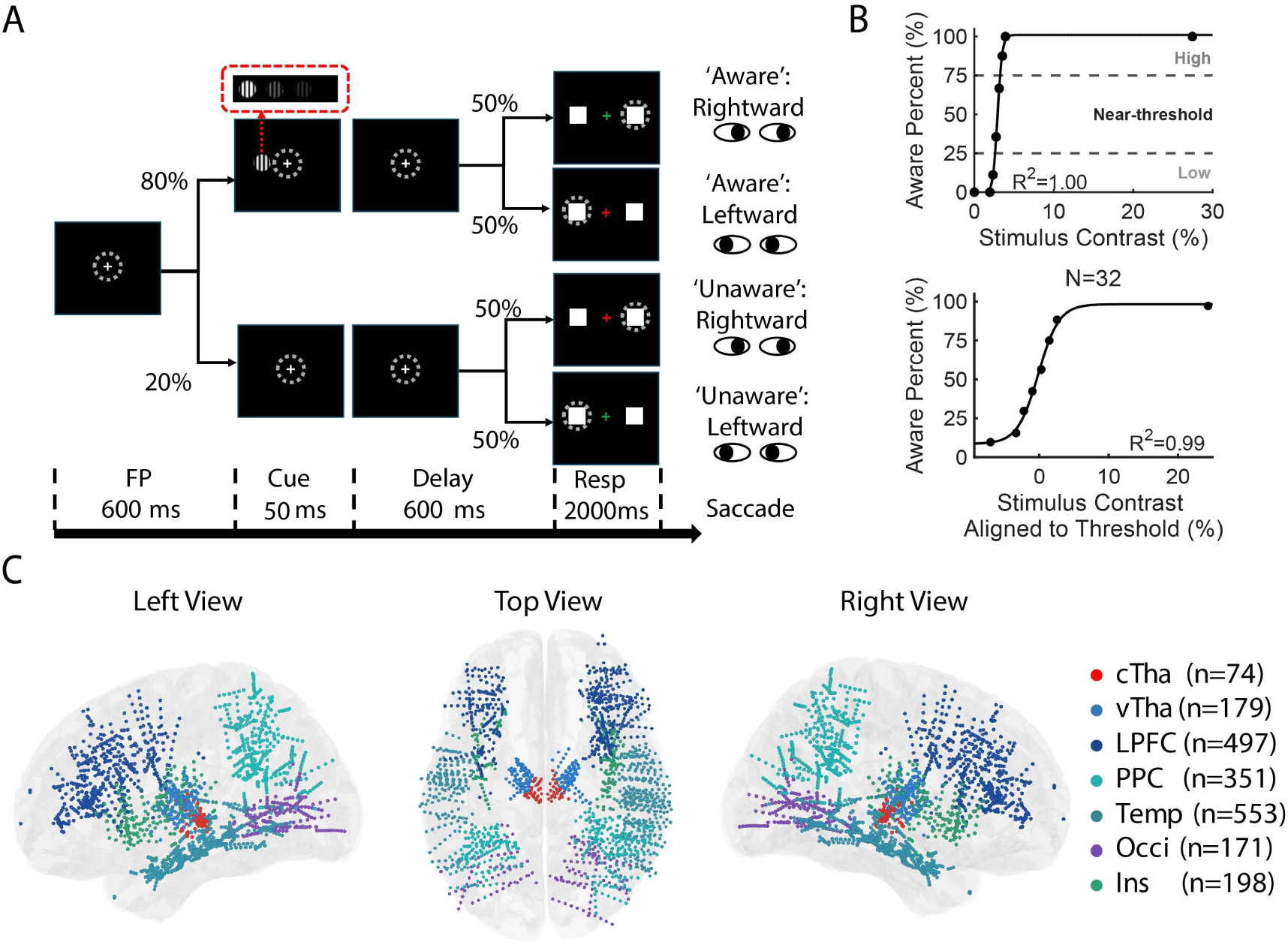
Visual awareness task, psychometric functions, and electrode localization. (A) Schematic illustration of the visual awareness task. (B) Psychometric detection functions from an example session (top) and across all subjects (bottom). Black dots indicate the percentage of conscious reports at each stimulus contrast, and black curves represent the fitted psychometric functions. Stimulus contrasts yielding 25–75% conscious reports were classified as near-threshold. For the population data (bottom), stimulus contrasts were aligned to each subject’s perceptual threshold (50% conscious reports). N, number of subjects; R², coefficient of determination. (C) Left, right, and top views of recording sites projected onto the Montreal Neurological Institute (MNI) brain template. Different colors indicate recording sites in different brain regions.

### sEEG results

During the task, we recorded sEEG activity from the the central thalamus (cTha) and other 6 brain regions (n=2023 recording sites) proposed to be important for conscious perception(*12, 26–32*): ventral thalamus (vTha), lateral prefrontal cortex (LPFC), posterior parietal cortex (PPC), middle and inferior temporal lobes (Temp), occipital lobe (Occi), Insula (Ins) (see Methods for details). Figure 1C shows the electrode locations in all seven brain regions (projected onto the Montreal Neurological Institute (MNI) ICBM152 brain template).

Because grating contrast was matched between conscious and unconscious trials, the differences of the post-stimulus activity during the delay period reflected consciousness-related activity, rather than simple visual responses to the appearance of grating. We examined the event-related potential activity at the recorded sites and totally yielded 600 consciousness-related sites for further analysis (Fig. 1, see Methods for details), i.e. 59/74 (80%) for cTha, 74/179(41%) for vTha, 141/497(28%) for LPFC, 114/351(32%) for PPC, 103/553 (19%) for Temp, 76/171(44%) for Occi, and 33/198(17%) for Ins, respectively. Nearly all (99.33%, 596/600) latencies of the identified consciousness-related activity were longer than 200 ms (Fig. S1), consistent with previous studies(*11, 14*).

### cTha showed early ERP activity immediately preceding conscious processing

We first examined whether there are early effects (0 - 200 ms) at the identified consciousness-related recording sites in cTha. Fig. 2A shows example sites which exhibited a sustained early activity within the first 200 ms after stimulus onset, preceding the larger subsequent evoked consciousness-related response after 200 ms. Although much weaker than the evoked activity, the early activity tended to be stronger in conscious than in unconscious trials and was relatively consistent across subjects, although the difference did not reach statistical significance in most subjects (Fig S2, p=0.063, 0.072, 0.075, 0.001, 0.065, 0.056 for sub 01-06, respectively). This early activity was also prominent at the population level at cTha, but being absent in vTha, LPFC, PPC, Temp, Occi and Ins (Fig. 2B). Direct comparisons of early consciousness-related activity across brain regions showed that, the early activity was significantly stronger in cTha than in the other five regions except Ins (Fig. 2C and Fig. S3), although only the comparison between cTha and LPFC survived the correction for multiple comparisons because of the weak effect. The other regions showed no significant effect (except vTha versus LPFC).

**Fig. 2.**
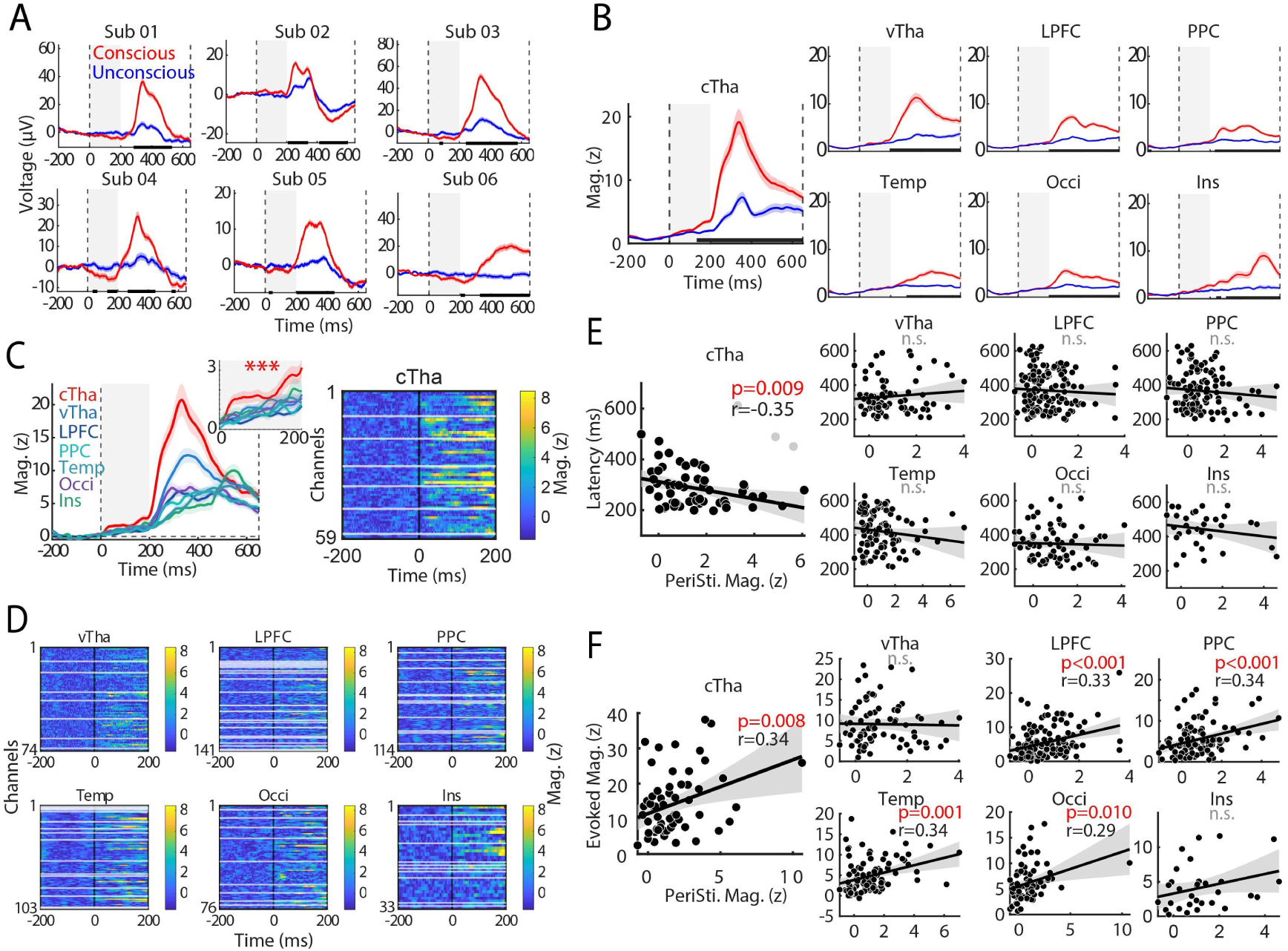
Characteristics of early consciousness-related activity in cTha. (A) Example sEEG activity from one cTha recording site per subject. Each panel shows the trial-averaged sEEG responses in conscious and unconscious conditions, with shaded areas indicating the standard error of the mean (SEM). Black dashed lines at 0 and 650 ms indicate grating onset and fixation-point color change, respectively. Thick black lines indicate significant differences between conditions (p < 0.01, FDR-corrected, independent-samples t-tests). Grey shading highlights the early consciousness-related activity (0–200 ms) in cTha. (B) Population sEEG activity of cTha and other brain regions. The figure format is the same as panel A but for population activity. (C) Comparison of consciousness-related ERP activity across brain regions. Left panel shows the consciousness-related activity in cTha and other brain regions. Activity in 0-200ms period was enlarged for clarity. Right panel shows heatmap of consciousness-related ERP activities across individual cTha recording sites. (D) Heatmaps of consciousness-related ERP activity across individual recording sites in other six brain regions. The figure format is the same as panel C but for other 6 brain regions. (E) Correlation between the strength of early activity (0–200 ms) and the latency of subsequent consciousness-related activity (200–650 ms). r, correlation coefficient; p, p value; n.s., not significant. (F) Correlation between the strength of early thalamic activity and subsequent consciousness-related activity.

We further explored whether the early activity shaped subsequent evoked consciousness-related activity. At cTha sites, the magnitude of early activity correlated negatively (r = −0.35, p = 0.009) with the latency of subsequent evoked consciousness-related activity (200 - 650ms), i.e. sites with stronger early activity tended to exhibit an earlier onset of evoked consciousness-related activity. This relationship was absent in the other regions. The magnitude of early activity also correlated positively (r = 0.34, p = 0.008) with the magnitude of the evoked consciousness-related activity, i.e. sites with stronger early activity tended to exhibit a stronger evoked consciousness-related activity. Together, these results suggest that early cTha activity shapes subsequent evoked consciousness-related activity.

### Early ERP activity in cTha showed inverse relationship with stimulus intensity

We asked whether the early component of thalamic activity was extrinsically evoked by the visual stimulus, or reflected intrinsic fluctuation. The assumption is that, the intrinsic activity would show inverse pattern with the late extrinsic one which were typically increase with stimulus strength, because weak stimulus require greater intrinsic compensation to reach the threshold for further conscious processing. We thus divided the conscious trials into three groups according to their contrast strength and compared the early activity among them. Fig. 3A shows the inverse ordering pattern of early activity with increasing stimulus strength at an example site in cTha (left, group averaged ERP; right, single-trial heatmap) i.e. early activity was stronger in the high contrast group than that in the low contrast group. At the population level, this phenomenon is also significant during the 100 to 200 ms period (p=0.013, High vs Low, Wilcoxon signed-rank test, uncorrected), showing the opposite pattern with the subsequent evoked activity (Fig. 3B-C), although the effect not survive the correction for multiple comparison across groups (p=0.133, Kruskal-Wallis test) due to the slight effect. Such pattern of early activity was absent in unconscious trials (Fig. 3D) and in the other six regions (Fig 3E).

**Fig. 3.**
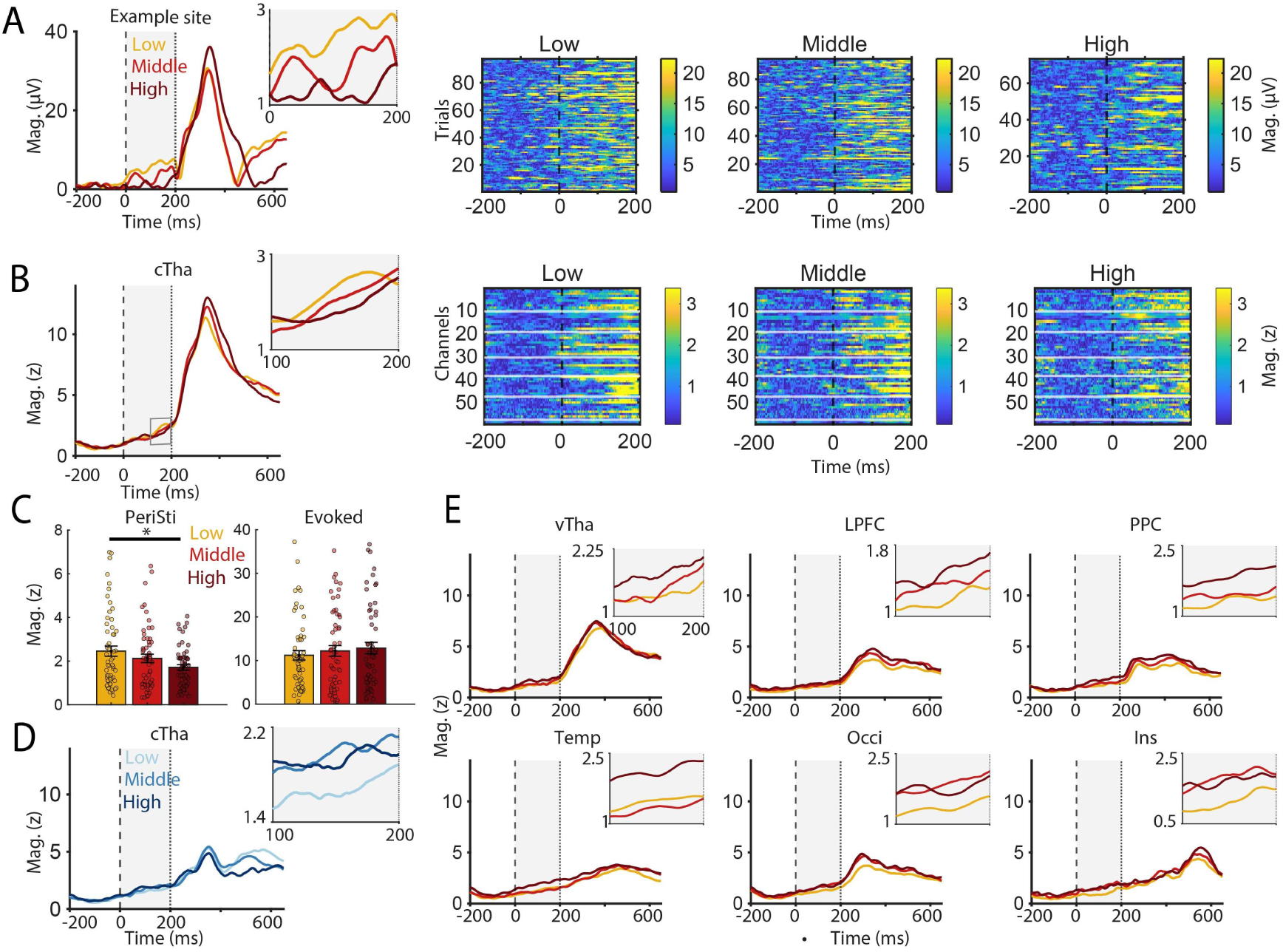
Inverse relationship between early ERP activity and stimulus contrast in cTha. (A) Example ERP responses across stimulus contrast levels at a representative cTha site during conscious conditions. Orange, red, and dark red lines indicate low, middle, and high stimulus contrasts, respectively. The early period (0–200 ms) is enlarged for clarity. (B) Population ERP responses across stimulus contrast levels in cTha during conscious conditions, displayed as in (A). (C) Comparison of ERP magnitudes across stimulus contrast levels in cTha during conscious conditions, separately for the early (0–200 ms, left) and late (200–650 ms, right) periods. (D) Population ERP responses across stimulus contrast levels in cTha during unconscious conditions, displayed as in (B). (E) Population ERP responses across stimulus contrast levels in six other brain regions during conscious conditions, displayed as in (B).

### cTha showed early phase concentration preceding conscious processing

We next examined whether the early cTha ERP was accompanied by changes in phase organization using inter-trial phase coherence (ITC). To exclude the contamination from evoked activity during signal filtering, the data was truncated at 200ms. Compared with unconscious trials, conscious trials showed stronger phase concentration in theta-band (2-8 Hz) at early period (Fig. 4A). Further analysis showed that during the early period, i.e. at approximately 100, 150, and 200 ms, cTha theta phases were more concentrated near the descending phase of the oscillation in conscious than in unconscious trials. (Fig. 4A, lower panels). Such phase concentration was consistent across all thalamic sites (Fig. S4A). Early consciousness-related phase concentration was also observed in alpha band but not in the other bands (Fig. S4B).

**Fig. 4.**
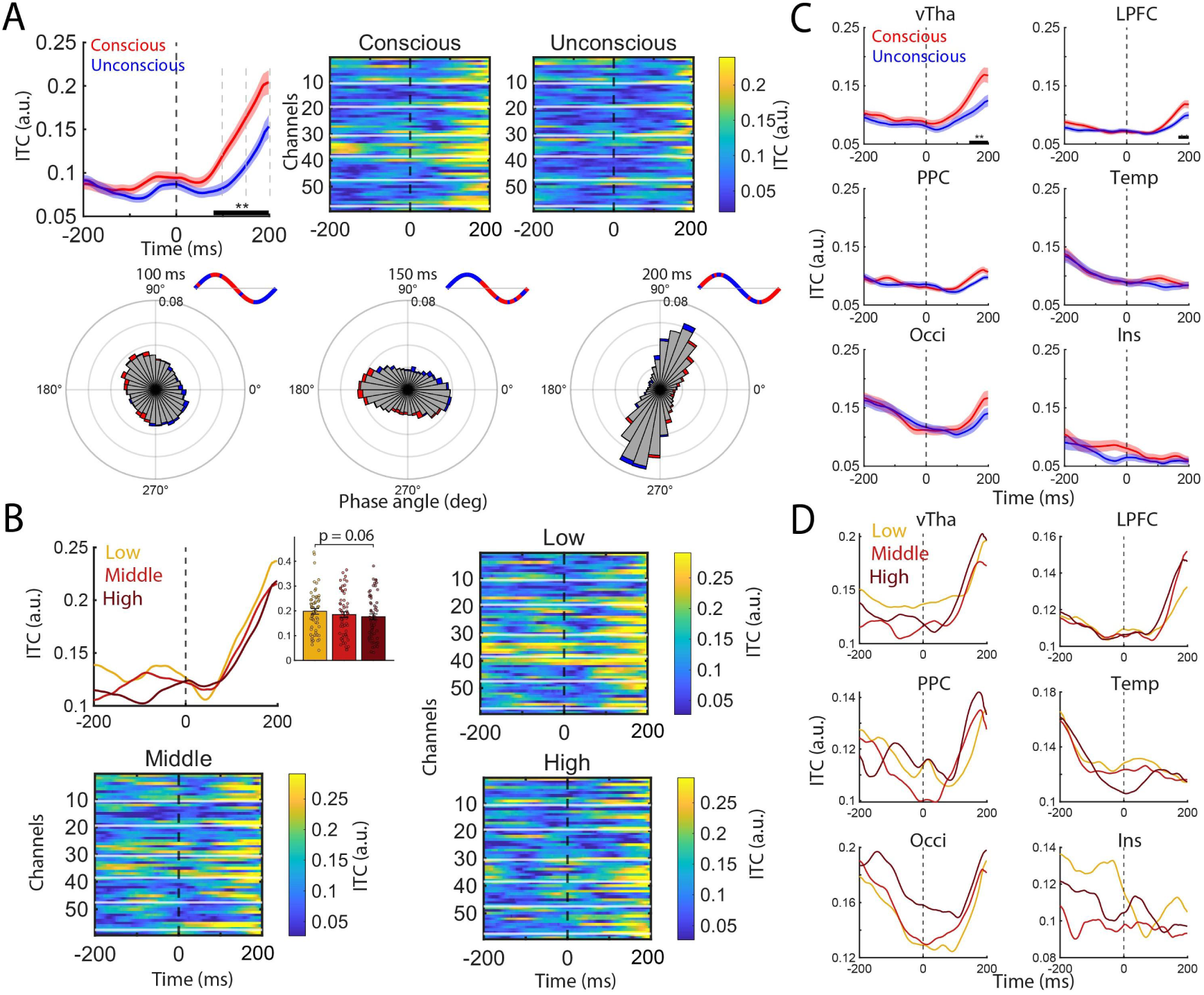
Early theta phase alignment in cTha preceding conscious processing. (A) Theta-band ITC(2–8 Hz) in cTha during conscious (red) and unconscious (blue) trials. Top, population-averaged ITC time courses and heatmaps across individual recording sites. Bottom, phase distributions at approximately 100, 150, and 200 ms. In the polar plots, the radius indicates probability and the angle indicates the phase bin; adjacent sinusoidal schematics illustrate phase position. (B) Theta-band ITC across low-, medle-, and high-stimulus contrast groups in cTha during conscious trials. Group-averaged ITC (left) and site-level heatmaps (right and bottom) are shown. (C) Theta-band ITC during conscious (red) and unconscious (blue) trials in the other six brain regions, displayed as in (A). (D) Theta-band ITC across low-, meddle-, and high-stimulus contrast groups in cTha during unconscious trials, displayed as in (B).

Similar to the ERP results, theta ITC during the early period also showed an inverse ordering trend across stimulus contrasts in conscious trials:, i.e. ITC_Low contrast_ > ITC_midlle contrast_ > ITC_high contrast_ (Fig 4B). Though not reach the statistical significance level (p=0.06, High vs Low, Wilcoxon sign rank test), it is consistent and evident in the individual-channel level (Fig 4B, right and bottom panels). vTha and LPFC also showed significant early consciousness-related ITC changes, but, similar with the other 4 regions, they did not exhibit an inverse ordering pattern (Fig. 4C-D). Such inverse ordering pattern was also absent in the other frequency bands (Fig. S4C).

Thus, early cTha activity also involved enhanced cross-trial phase alignment in the theta band.

### cTha showed early connectivity enhancement preceding conscious processing

We next explored that whether early functional coupling between cTha and other regions contribute to subsequent conscious processing using phase-locking value (PLV). The local PLV at alpha band within cTha showed an early consciousness-related increase (Fig. 5A), which was absent in the coupling with other regions (Fig. 5B) or in other bands (Fig. S5). Importantly, the early PLV strength within cTha was positively correlated with the strength of subsequent evoked activity (Fig. 5C left panel, r = 0.48, p < 0.001), i.e. greater local phase locking was thus associated with stronger evoked activity, linking early cTha coupling to the magnitude of the subsequent response. This correlation was also significant, though weaker, for the cTha-vTha pair (Fig. 5C, right panel; r = 0.25, p= 0.005) but was absent in other five cTha-seeded regional pairs (Fig S6).

**Fig. 5.**
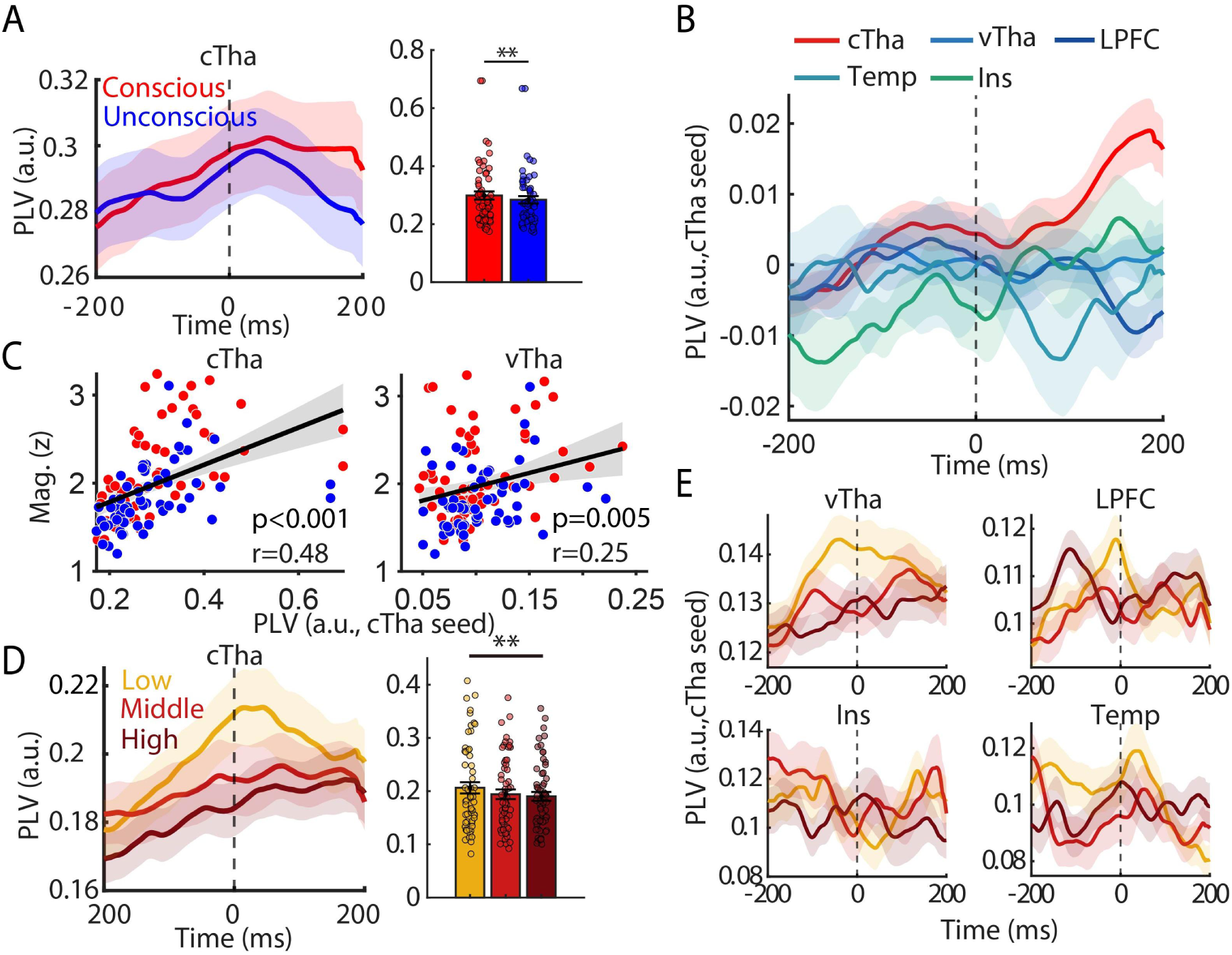
Early connectivity enhancement preceding conscious processing. (A). Within-cTha alpha-band PLV (8–12 Hz) for conscious (red) and unconscious (blue) trials. Left panel, population averaged curve; right panel, comparison between the mean PLV in 0-200ms under two conditions. (B) Comparison of consciousness-related early PLVs (cTha seeds) across all regions. Colors indicate different brain regions pairs. (C) Correlation between early PLV and subsequent response magnitude for within-cTha (left) and cTha–vTha (right) coupling. Red and blue dots represent observations from conscious and unconscious trials, respectively. Black lines show pooled linear fits and gray bands indicate the 95% confidence intervals; r denotes Pearson correlation. (D) Population PLV across stimulus contrast levels in cTha during the conscious condition. Orange, red, and dark red lines indicate low, middle, and high stimulus contrasts, respectively. (E) Population PLV across stimulus contrast levels in the other six brain regions during the conscious condition, displayed as in (D).

Moreover, within-cTha PLV also showed an inverse ordering trend across stimulus contrasts during the early period (100 - 200 ms) in conscious trials (Fig. 5D, Low versus High, p = 0.0048, Wilcoxon signed-rank test, uncorrected; mean PLVs 0.200, 0.193, and 0.190 for the Low, Middle, and High groups, respectively). This ordering therefore constituted a graded trend consistent with the early ERP and phase concentration patterns. This pattern appeared exclusively in within-cTha PLV, but not the PLVs between cTha and other regions (Fig. 5E).

### cTha showed a sustained shift in critical dynamics preceding conscious processing

Finally, we asked whether the early thalamic effects could be explained by the collective dynamical state, i.e. criticality, of cTha. We used the branching ratio (BR)(*33*) as a probe and estimated the BR changes from -500 to 600 ms by conducting the sliding-window analysis (window length 800ms, step size 100ms) in conscious and unconscious trials. In conscious trials, cTha (N=7 subjects) exhibited a significantly higher BR (p < 0.05 uncorrected, Wilcoxon sign-rank test) which sustained from - 300ms to 200ms, compared to unconscious trials (Fig. 6A). And the higher BR in conscious trials corresponded to a longer estimated autocorrelation time. This means that, conscious perception is more likely to occur when thalamic activity is closer to the near-critical regime (BR=1) and exhibits more persistent temporal correlations during the peristimulus period.

**Fig. 6.**
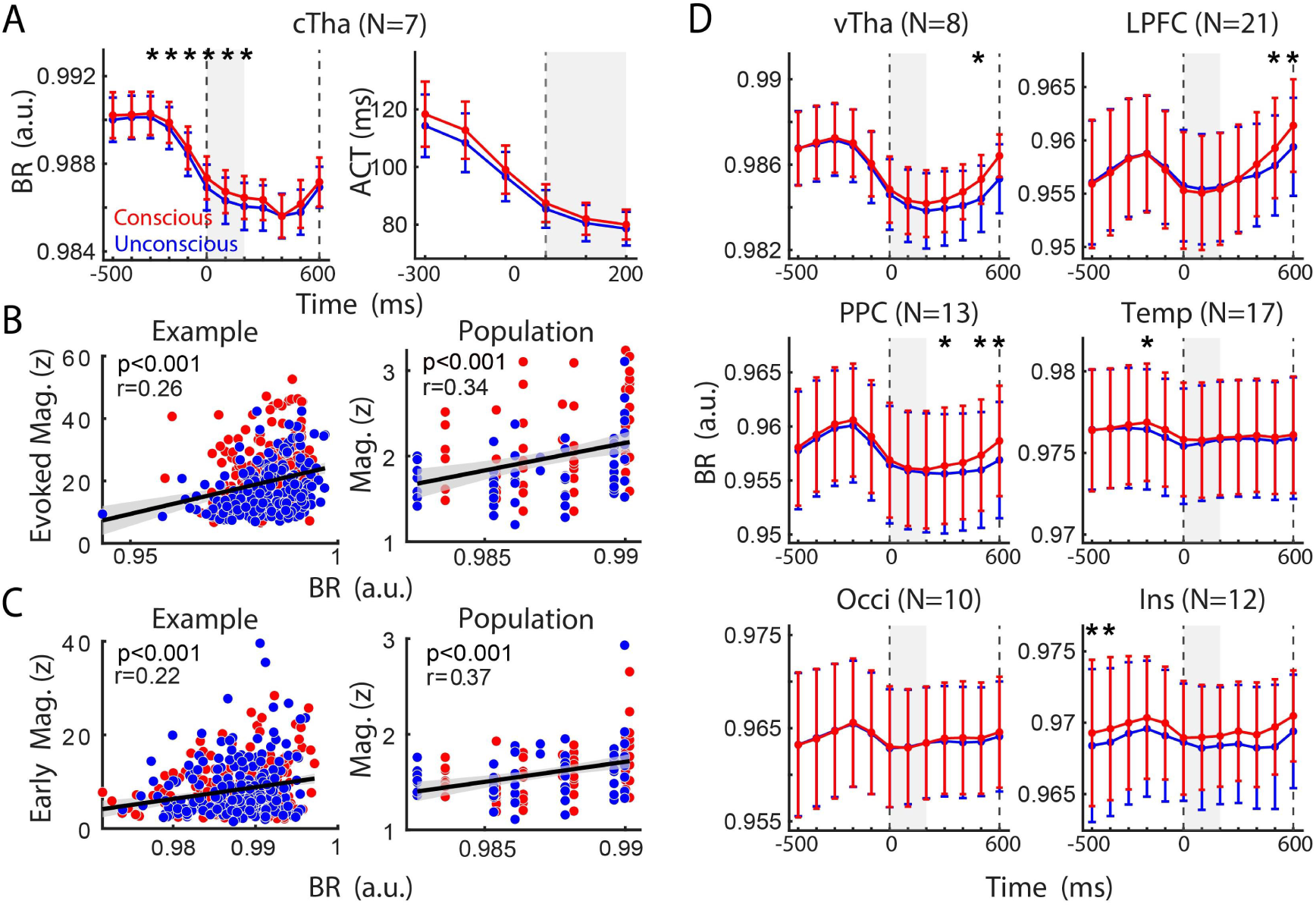
Peristimulus BR and autocorrelation time. (A) BR (left) and autocorrelation time (ACT; right) in cTha during conscious (red) and unconscious (blue) trials. Estimates were obtained using 800-ms sliding windows with 100-ms steps. Time points indicate window endpoints (e.g., 0 ms corresponds to −800 to 0 ms). Asterisks indicate significant differences between conditions (p < 0.05, uncorrected, two-sided paired Wilcoxon signed-rank tests across subjects). (B) Correlations between prestimulus BR (−800 to 0 ms) and the magnitude of subsequent stimulus-evoked ERP activity (200–650 ms) at a representative recording site (left) and at the population level (right). (C) Correlations between prestimulus BR (−800 to 0 ms) and the magnitude of early ERP activity (0-200 ms), displayed as in (B). (D) BR (left) and ACT (right) during conscious (red) and unconscious (blue) trials in six other brain regions, displayed as in (A).

Criticality was correlated with local neural activity. In cTha, BR at stimulus onset was correlated positively with the magnitude of the evoked response (r = 0.34, p = 0.0002; Fig. 6B) and also with early activity (r = 0.37, p < 0.001; Fig. 6C). Representative sites showed correlations in the same direction (Fig. 6B, evoked activity: r = 0.26, p < 0.001; Fig. 6C, early activity: r = 0.22, p < 0.001). These findings suggest that an early shift of cTha dynamics toward a more critical regime and provides a favorable dynamical state for the subsequent large-amplitude evoked response.

Other brain regions, including vTha (N = 8), LPFC (N = 21), PPC (N = 13), Ins (N = 12), Occi (N = 10), and Temp (N = 17), did not show sustained differences in BR during the peristimulus period, although some cortical regions showed differences in later or isolated time windows. Thus, the early shift in critical dynamics associated with conscious perception was selective for cTha.

## Discussion

Why the same sensory input sometimes reaches consciousness and sometimes fails to do so is a fundamental question in the neuroscience of consciousness. By recording sEEG activity from the thalamus and other consciousness-related regions while patients performed a conscious perception task, we found that momentary intrinsic fluctuations in high-order thalamus, including early ERP activity, phase concentration, and local functional connectivity enhancement, as well as the shift in the collective dynamical state toward critical regime, shape whether the thalamic gate opens during conscious perception. These results suggest that intrinsic fluctuations in high-order thalamus directly shape how the thalamus gates conscious processing and thus modulate the probabilistic emergence of conscious perception.

### Early consciousness-related activity appeared exclusively in high-order thalamus but not in other consciousness-related regions

A striking feature of the present results is the anatomical selectivity of the early consciousness-related activity. Within the first 200 ms after stimulus onset, several robust early activities emerged in the high-order thalamus preceding conscious processing, including the ERP activity, phase concentration and within-region phase coupling, whereas no comparable early effects were detected in the other examined consciousness-related regions, namely vTha, LPFC, PPC, the middle and inferior temporal lobes, occipital lobe and the insula. This spatiotemporal specificity places the high-order thalamus at a uniquely early stage of the neural processes leading to conscious perception. Rather than reflecting a kind of widespread conscious processing, the early thalamic signal may therefore represent an initiating or enabling process that precedes the recruitment of distributed cortical networks. This interpretation is further supported by the observation that the magnitude of the early thalamic response predicted both the latency and amplitude of the subsequent consciousness-related activity, suggesting that the early thalamic state is functionally linked to the later development of the conscious processing.

More broadly, these findings argue against the view that consciousness-related activity first emerges within sensory or association cortex(*27, 34, 35*) and is subsequently relayed to the thalamus. Instead, they support a hierarchical temporal architecture in which the high-order thalamus occupies an upstream position, potentially establishing the prerequisites under which later cortical and thalamocortical activity can develop into conscious processing.

### Early consciousness-related activities primarily reflect the intrinsic state

Notably, among consciously perceived trials, these early activities scaled inversely with stimulus strength, in striking contrast to the conventional positive stimulus-response relationship, e.g. observed in the later evoked consciousness-related activity (Fig. 3B). This suggests that intrinsic activity complements stimulus-evoked activity at the threshold for conscious processing, i.e. during successful conscious perception, lower-contrast stimulus trials were more likely to be associated with stronger intrinsic activity that helped reach the thalamic threshold for conscious processing. Moreover, these early activity patterns were not apparent in other brain regions or in unconscious trials, indicating the unique importance of intrinsic fluctuations in high-order thalamic nuclei during the probabilistic emergence of conscious perception.

Together, these findings suggest that early thalamic activity primarily reflects intrinsic fluctuations. The complementary relationship between intrinsic and extrinsic activity in high-order thalamus provides a simple but plausible explanation for the moment-to-moment variability of conscious perception of near-threshold stimuli.

### Early thalamic activity directly links to subsequent evoked consciousness-related activity and shapes conscious perception

Importantly, this intrinsically arising early thalamic activity is not an isolated background fluctuation; rather, it emerges transiently at approximately 0-150ms and sustained to 200m when the stimulus-evoked response arises. Thus, intrinsic thalamic activity directly interacts with the incoming stimulus-evoked response and collaborated in a complementary manner. Beyond this, intrinsic activity further shapes subsequent conscious processing and perceptual outcomes. Across high-order thalamic sites, stronger early activity was associated with a shorter latency and a greater magnitude of the later consciousness-related response (Fig.2E-F). This relationship was further supported at the oscillatory level: early phase organization at low-frequency, including phase concentration and synchronization (Figs. 4–5), predicted the magnitude of the later evoked response, suggesting that the instantaneous temporal organization of intrinsic thalamic activity influences the incoming conscious processing.

Moreover, early phase coupling increased primarily within high-order thalamus rather than between thalamus and other examined cortical regions (Fig. 5A), suggesting that the earliest conscious processing may not require an already established large-scale thalamocortical network state. Instead, local thalamic populations may first enter a more coordinated configuration that changes their susceptibility to incoming input.

### Early thalamic activity might reflect intrinsic fluctuations in thalamic criticality

The convergence of several types of early activity suggested a changing thalamic state. To elucidate the underlying dynamical state principle, we focused on criticality, which has been shown to correlate closely with global conscious states and to enable optimal neural computation(*20, 22, 36, 37*). The criticality, indexed by BR, difference between conscious and unconscious trials sustained since 300ms before the stimulus onset to 200ms afterward, suggesting that the change in criticality was primarily intrinsic rather than stimulus-evoked. During conscious trials, activity in high-order thalamus was closer to the critical regime (BR=1) compared to unconscious trials. This can be understood as follows: during unconscious trials, a weak sensory perturbation arriving in a relatively subcritical thalamic state may therefore dissipate before it can effectively recruit downstream circuits; by contrast, the same perturbation arriving when thalamic dynamics lie closer to criticality would be more likely to persist and reverberate, increasing the probability of subsequent conscious processing. This criticality difference diminished at approximately 200ms, when the evoked activity emerged, further supporting this proposal.

To the best of our knowledge, this result provides the first evidence that fluctuations in the critical state of high-order thalamus shape probabilistic conscious perception. Conscious states and conscious contents can therefore be unified within a single quantitative dynamical framework: conscious states correspond to the system’s position in phase space (subcritical/critical/supercritical)(*20, 21, 38*), whereas conscious content corresponds to transient phase transitions triggered by specific sensory inputs near the critical state.

### Intrinsic fluctuations help to explain why the thalamic gate for conscious perception opens on some occasions but not on others

The present study extends our previous finding(*16*) that high-order thalamic nuclei gate conscious perception through interactions with the prefrontal cortex and further addresses why the thalamic gate opens on some trials but not others. Intrinsic fluctuations in high-order thalamus may first determine the gain applied to an incoming sensory event; then the processed activity that crosses the thalamic threshold for conscious processing would be sufficiently amplified and recruits thalamocortical interactions and distributed cortical processing, leading to successful conscious perception; By contrast, the activity without enough gain and fails to cross the thalamic threshold will dissipate shortly, leading to unconscious perceptual outcome. In this framework, the high-order thalamus is not merely a downstream participant in conscious processing; rather, it may initiate the conscious processing and function as a state-sensitive control system situated at the interface between ongoing brain dynamics and externally driven computation. Such a mechanism is consistent with the broad anatomical and physiological organization of higher-order thalamic nuclei.

This interpretation also provides broader implications for understanding various cognitive functions. Plenty of cognitive functions, e.g. working memory, attention etc, are modulated by intrinsic brain states(*39*). If higher-order thalamic dynamics regulate the susceptibility of distributed cortical circuits more generally, moment-to-moment fluctuations in high-order thalamus could provide a common mechanism through which internal brain state directly shapes multiple forms of cognition. Testing how intrinsic state – extrinsic inputs interact in high-order thalamus across different cognitive domains will be important for establishing a general mechanism by which how brain intrinsic states shape information processing and further cognition.

### The observed thalamic fluctuations are unlikely to be explained by arousal fluctuations or attentional sampling

Several findings in our data argue against attentional sampling or arousal states as sufficient explanations for the thalamic effects. First of all, the early thalamic activities were transient, emerging around stimulus onset and lasting approximately 100-200 ms. This differs from the reported fluctuations in arousal states which typically last at least several seconds (*40, 41*). Second, in our paradigm, the location and timing of the grating were fixed across trials for each subject, minimizing the differences in attentional allocation between conscious and unconscious trials. The results further showed that the early consciousness-related activity was selective for the high-order thalamus but largely absent from LPFC and PPC, the key regions involved in attentional control(*42, 43*). Early phase coordination also increased primarily within high-order thalamus rather than across a broader frontoparietal network. Together, these results suggest that the observed thalamic fluctuations are unlikely to be explained solely by arousal fluctuations or attentional sampling.

## Methods

### Data acquisition

Data acquisition procedures followed those described previously (*16, 24, 25*). Briefly, 32 adult patients (26 males; mean age ± SD: 29.13± 8.8 years) with drug-resistant epilepsy or headaches were recruited from the Chinese PLA General Hospital and the General Hospital of Western Theater Command. All patients had normal or corrected-to-normal vision. Depth electrodes (Sinovation Medical Technology, Beijing, China), each containing 8-20 contacts (0.8 mm in diameter, 2 mm in length, and 1.5 mm intercontact spacing), were implanted solely based on clinical requirements. sEEG signals were referenced to a white-matter contact, sampled at 1 kHz, bandpass-filtered at 1–250 Hz, and notch-filtered at 50 Hz using Neuracle or Nicolet recording systems. Stimulus-triggered electrical pulses were simultaneously recorded to synchronize stimulus onset with neural activity.

Experiments were conducted in a quiet, dimly environment. Visual stimuli were presented on a 24-inch LED monitor (Admiral Overseas Corporation; 144 Hz, 1,920 × 1,080 pixels), using MATLAB (MathWorks) and Psychtoolbox-3 (PTB-3; Brainard and Pelli). Eye movements were monitored at 1 kHz using an infrared eye tracker (EM2000C, Beijing Jasmine Science & Technology). Patients continued taking their prescribed medications during experiments. All subjects provided informed consent, and the study was approved by the Ethics Committee of the Chinese PLA General Hospital (S2022-457-01, S2023-809-02).

### Electrode localization

Electrode locations were determined by coregistering the postoperative computed tomography (CT) scans with the preoperative T1-weighted MR images using a standardized mutual information method in SPM12 (*44*). Individual cortical reconstruction and parcellation were performed using FreeSurfer (*45*)with the Desikan– Killiany atlas, while thalamic nuclei were segmented using the FreeSurfer thalamic atlas (*46*). Recording sites were anatomically labeled based on individual cortical and thalamic parcellations using the automated electrode labeling function in Brainstorm. Montreal Neurological Institute (MNI) coordinates were obtained through nonlinear registration to the MNI ICBM152 template using SPM12.

Corresponding the Freesurfer atlas, the selected brain regions were defined as follows: cTha included CM (central median nucleus), MDm (mediodorsal medial magnocellular nucleus), Pf (parafascicular nucleus), CeM, MDl (mediodorsal lateral parvocellular nucleus); vTha included VA (ventral anterior nucleus), VLa (ventral lateral anterior nucleus), VLp (ventral lateral posterior nucleus), VPL (ventral posterolateral nucleus); LPFC included rostralmiddlefrontal, caudalmiddlefrontal, parsopercularis, parstriangularis; Occi included lateraloccipital, cuneus, pericalcarine, lingual; Temp includes inferiortemporal, middletemporal; PPC included superiorparietal, inferiorparietal; Ins included insula.

### Experimental task

As illustrated in Fig. 1, a trial started when a fixation point (0.5° × 0.5°, white cross) appeared at the center of the screen (radius of the eye position check window = 4°, shown as the dotted circle). After the subject fixated on the white cross for 600 ms, a cue stimulus (Gabor grating, 2 × 2° circle) was presented for 50 ms at a fixed position (7°) on the left (or right) side of the fixation point. The grating stimulus was presented contralateral to the major implanted hemisphere of electrodes. In 70% of the trials, the grating contrast (Weber contrast) was maintained near the subject’s perceptual threshold by a 1 up/1 down staircase method, and the step was 0.39% of the highest contrast of the screen. In 10% of the trials, the stimulus contrast was well above the threshold, and in the other 20% of the trials, the stimulus contrast was 0, namely, no stimulus appeared. After another 600 ms delay, the color of the fixation point turned red or green, and two saccade targets (1 × 1°, white square) appeared at fixed positions (10°) on the left and right sides of the fixation point. If the grating was seen, a green fixation point indicated that subjects should make a saccade to the right target, whereas a red fixation point indicated that subjects should make a saccade to the left target. If the grating was not visible, the rule of the saccadic direction was inverted. Suprathreshold and stimulus-absent trials served as controls to assess task comprehension and performance. Before data collection, subjects completed 1–2 training sessions, during which individual perceptual thresholds were determined. Each experimental session comprised 180 trials, with an intertrial interval of 800 ms. Each subject completed 5–7 sessions.

### Data analysis

Trials were excluded if subjects broke fixation (eye position outside the 4° × 4° fixation window for >100 ms) or failed to respond within the maximum response period after target onset (2,000 or 5,000 ms based on individual performance). More than 85% of trials were retained for subsequent analyses for each subject.

Within each session, trials were classified as conscious or unconscious based on subjects’ reports of grating visibility. Stimulus contrasts were further categorized into three levels according to the proportion of conscious reports: high (>75%), near-threshold (25–75%), and low (<25%). Given the limited number of low-contrast conscious and high-contrast unconscious trials, we focused on four conditions: low-contrast unconscious, near-threshold unconscious, near-threshold conscious, and high-contrast conscious. On average, these conditions comprised 142, 274, 274, and 238 trials per subject (15.31%, 29.52%, 29.52%, and 25.65%, respectively). Analyses primarily compared near-threshold conscious and unconscious trials to investigate the neural and behavioral correlates of conscious perception.

### Behavioral data analysis

#### Psychometric curve fitting

The psychometric curve was fitted via the following formula:

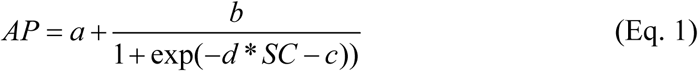

where AP indicates the ratio of reported ‘conscious’, SC indicates the stimulus contrast, and a-d indicate individual fitting parameters.

For population-level psychometric curve fitting, adjacent contrast levels were averaged, resulting in 8 contrast levels per session.

### sEEG data analysis

#### Data preprocessing

sEEG data were preprocessed using EEGLAB in MATLAB (*47*). Signals were bandpass-filtered at 1–250 Hz and notch-filtered at 50, 100, 150, 200, and 250 Hz using finite impulse response (FIR) filters. Data were epoched from −1,500 to 1,799 ms relative to grating onset. Noisy or artifactual recording sites were excluded based on visual inspection of event-related potentials (ERPs) and power spectral density (PSD). To minimize volume conduction and shared signals across recording sites, bipolar re-referencing was performed by subtracting signals from adjacent contacts on the same electrode. The anatomical location of each bipolar recording site was defined as the midpoint between the two corresponding contacts.

### Definition of visual consciousness-related activity at recording sites

To identify consciousness-related sites, we compared sEEG amplitudes between near-threshold conscious and unconscious trials at each time point from 0 to 650 ms after grating onset using independent-samples t-tests. Recording sites showing significant differences (p < 0.01, false discovery rate [FDR] corrected across time points) for more than 20 consecutive milliseconds were defined as consciousness-related sites. The earliest time point of sustained significant divergence between the conscious and unconscious trials was defined as the latency.

### Population ERP activity calculation

We normalized the ERP activity of each channel using z-score while calculating the population ERP activity:

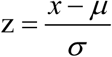

While µ and σ denote the mean and std standard deviation, respectively, of the ERP activity between -200 to 0 ms.

Given the complex physiological interpretation of sEEG signal polarity (*48*), we quantified consciousness-related activity based on the magnitude of the ERP difference between conscious and unconscious trials, irrespective of its polarity. Specifically, for each consciousness-related site (p < 0.01, FDR corrected), we calculated the absolute values of ERPs under conscious and unconscious conditions (or the ERP differences between them) and averaged these values across sites within brain regions to obtain population-level activity.

In the contrast-level analysis (e.g. Fig 3), trials were sorted in ascending order of stimulus contrast within each subject and perceptual condition (conscious and unconscious). For K trials, d = round(K/3); sorted trials 1 to d, d + 1 to 2d, and 2d + 1 to K formed the Low, Middle, and High groups, respectively. Thus, the groups contained approximately equal trial counts.

Moreover, because the latencies of evoked activity differed slightly across contrast groups and undermined the clarity of the preceding inverse ordering pattern, we realigned ERPs using the inflection time between early and evoked activity in the low contrast group as a reference (∼200 ms). Specifically, we shifted the population ERPs of the Low, Middle, High groups forward by 0, 15 and15 ms, respectively, and then performed further comparisons.

### Inter-trial coherence (ITC)

We focused on phase-based measures rather than spectral power, as power estimates can be sensitive to windowing effects. While calculating the band-limited ITC, we filtered the sEEG signals between -1500 to 200 ms and extracted the band-limited phase via the Hilbert transformation using Brainstorm Toolbox (*49*). For each recording site that showed consciousness-related ERP activity, ITC was calculated as:

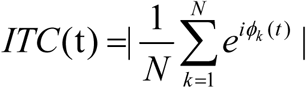

where N is the number of trials and φₖ(t) represents the instantaneous phase at time t in trial k. ITC values range from 0 (no phase consistency) to 1 (perfect phase consistency). ITC was then averaged across recording sites within each brain region. The ITC were subsequently averaged across the sites whitin brain regions. For polar histograms, phases were expressed on 0–360°, binned in 10° intervals, normalized to probabilities within each site, and averaged across sites.

### Phase-locking value (PLV)

The time-across PLV was calculated via the brainstorm toolbox (*49*). For a pair of filtered signals *x̃*(*t*) and *ỹ*(*t*) at -1500 to 200 ms:

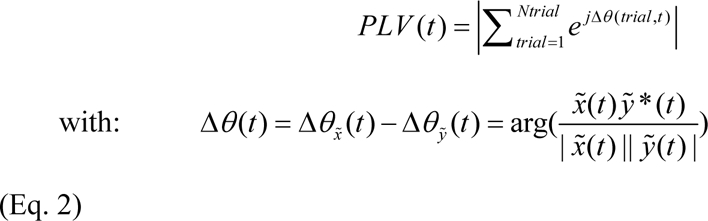

where Δθ(t) indicates the phase difference between two signals at time t.

### Branching ratio (BR) and autocorrelation time (ACT)

BR and autocorrelation time (ACT) were estimated using the multistep regression (MR) framework (*33, 50*), adapted to continuous sEEG signals. This novel estimator is suitable for analyzing the task-related sEEG recordings because it allows reliable estimation under subsampling. Briefly, for each trial and time window, neural activities from all recording sites within each brain region was used to estimate lag-dependent regression slopes (r_ₖ_) through linear regression:

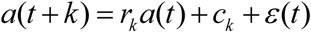

where a(t) represents neural activity at time t, k denotes the temporal lag, cₖ is the intercept, and ε(t) is the residual error. Regression slopes were calculated for lags of 1– 35 ms and subsequently fitted to an exponential model:

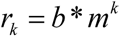

where b is the amplitude parameter and m represents the estimated BR. The ACT was then calculated as:

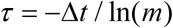

where Δt = 1 ms is the sampling interval. For 0 < m < 1, a higher BR corresponds to longer ACT, indicating a slower decay of temporal correlations and greater temporal persistence of neural activity.

To minimize stimulus-evoked contributions, the condition-averaged event-related potential (ERP) was subtracted from each trial at each recording site. BR and ACT were estimated using 800-ms sliding windows with a step size of 100 ms, yielding 12 windows with endpoints ranging from −500 to 600 ms relative to stimulus onset. Trial-level BR and ACT values were averaged according to analysis requirements. Group differences between conscious and unconscious conditions were assessed using paired Wilcoxon signed-rank tests across subjects.

## Acknowledgements

We thank the participants for volunteering to take part in the study.

## Funding

Ministry of Science and Technology of the People’s Republic of China grant Brain Science and Brain-like Intelligence Technology-National Science and Technology Major Project 2021ZD0204300;

National Natural Science Foundation of China grant 324B2031, 32595492;

National Key Research and Development Program of China grant 2025YFE0218700.

## Author contributions

Conceptualization: FZP, ZMS

Methodology: FZP, SHF, CX, YSX, ZJX, LZ

Software: FZP

Validation: FZP

Formal analysis: FZP, ZMS

Investigation: FZP, DYY

Resources: FZP, ZHL, ZMS, CL, SHF, CX, PAJ

Data curation: FZP, DYY, ZHL, ZMS

Writing – original draft: FZP, ZMS

Writing – review & editing: FZP, CL, SHF, ZMS

Visualization: FZP, ZMS

Supervision: FZP, ZMS

Project administration: FZP, ZMS, CL

Funding acquisition: ZMS, FZP, CL

## Competing interests

All the authors declare no competing interests.

## Data and materials availability

Electrophysiological data were analyzed using MATLAB in conjunction with the toolboxes mentioned above. Source data and customized MATLAB analysis scripts can be made publicly available according to the journal’s policy.

## Supplementary Materials

**Fig. S1.**
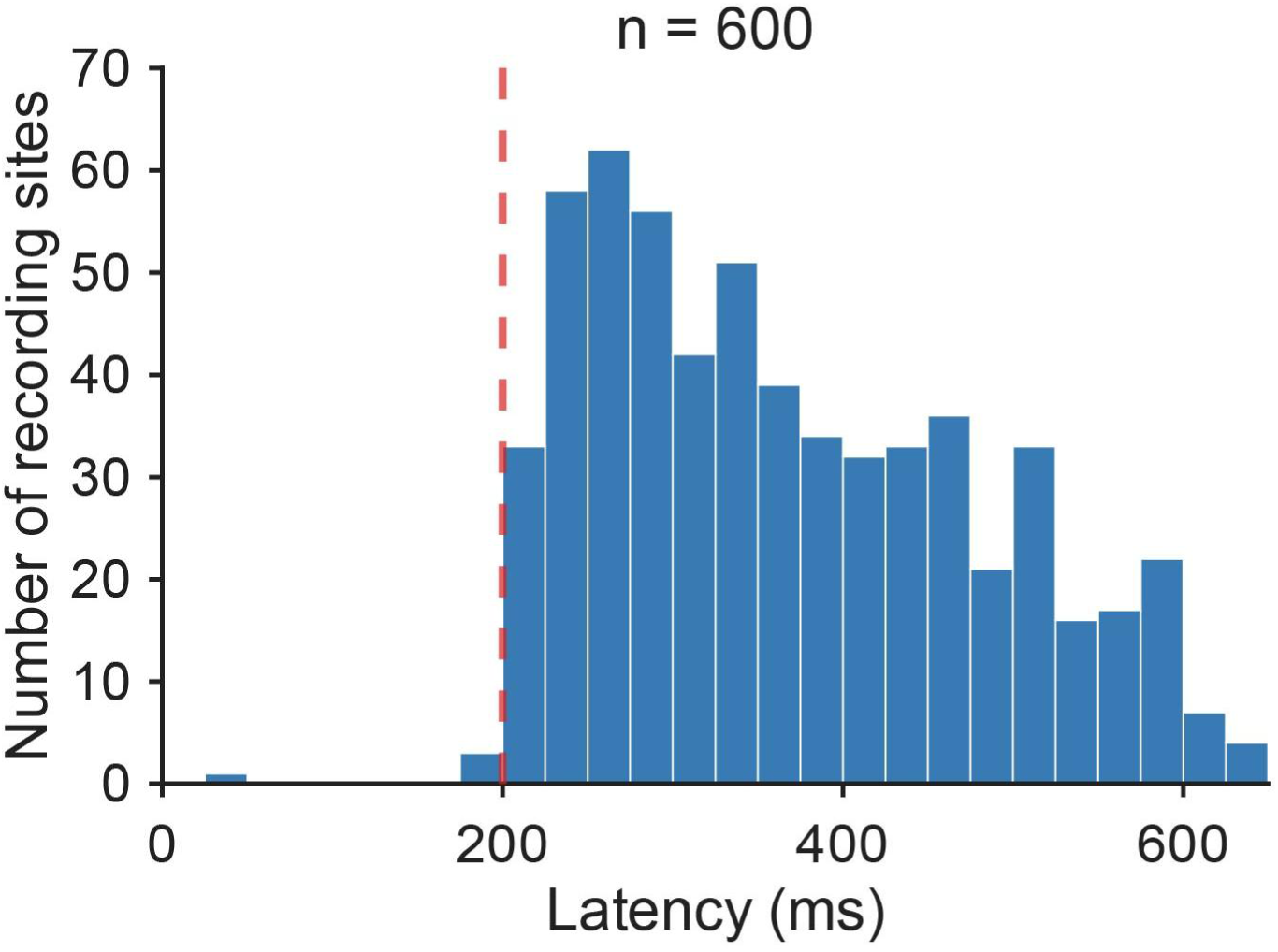
Distribution of the latencies of identified consciousness-related ERP activity.

**Fig. S2.**
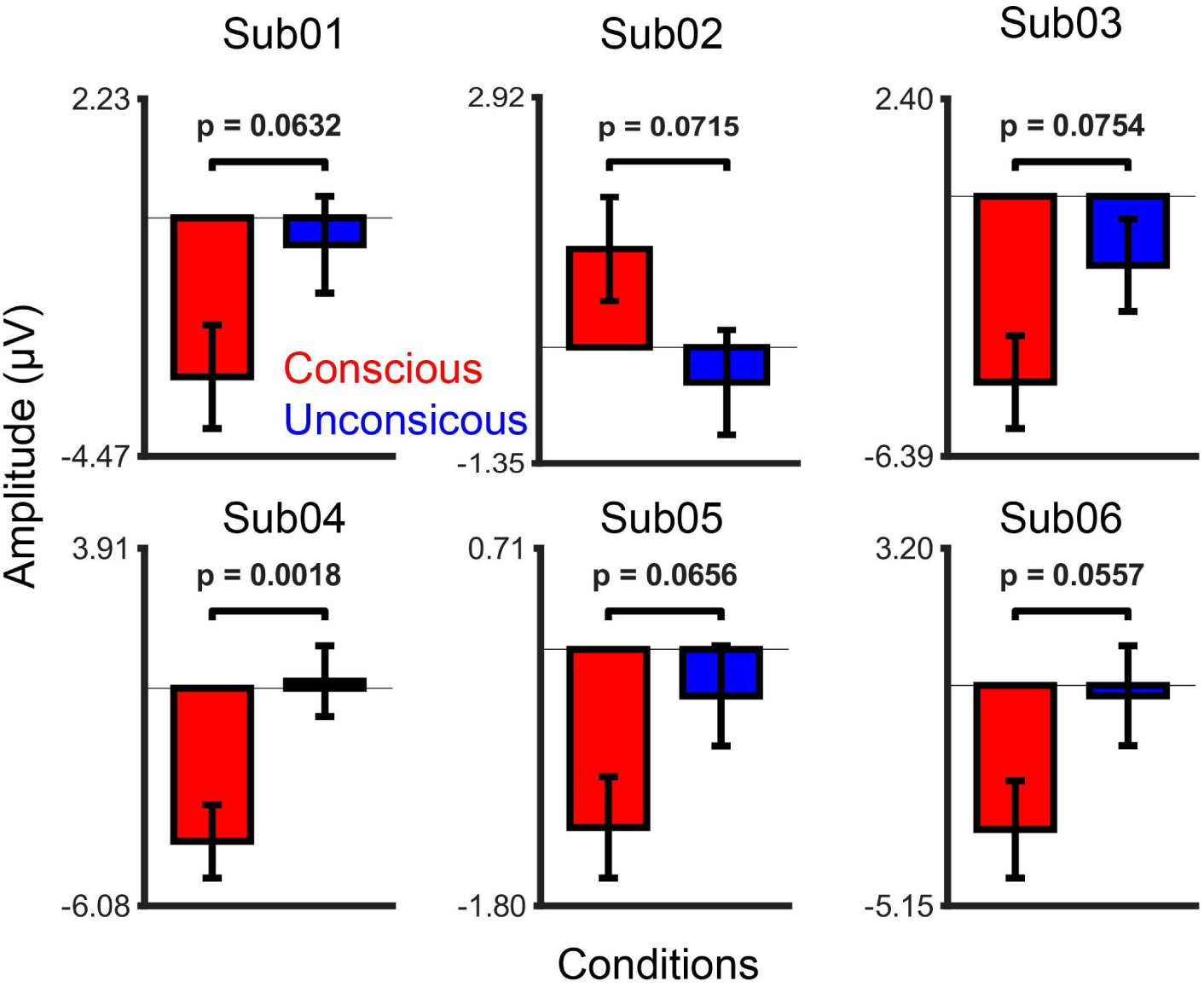
Statistical t tests of the mean amplitudes during 0-200ms in conscious and unconscious trials.

**Fig. S3.**
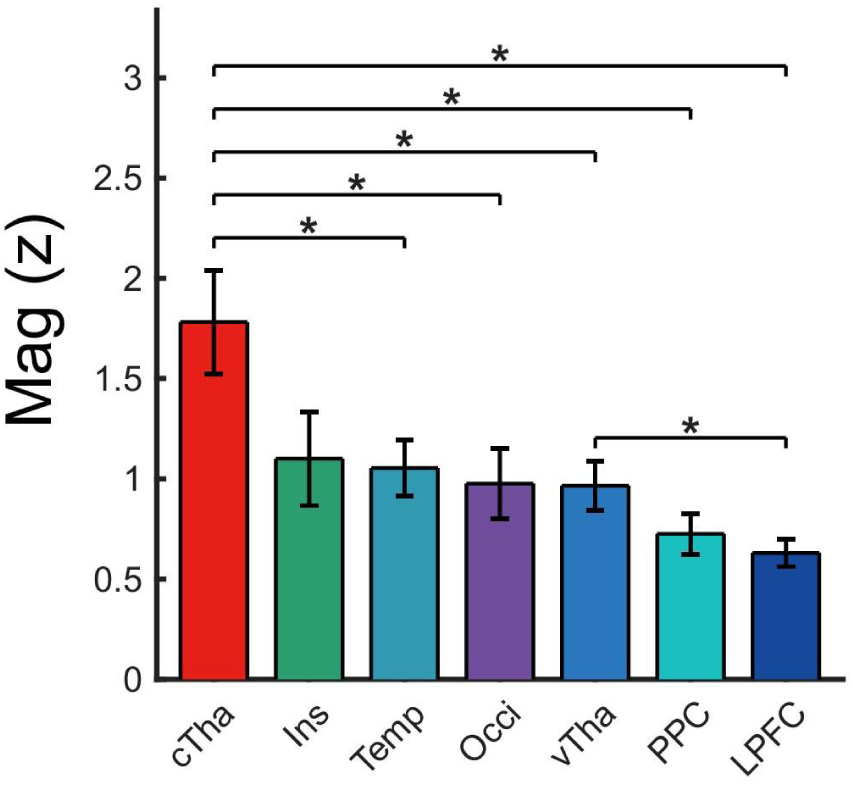
Comparison of consciousness-related ERP activity across brain regions. *, p<0.05 uncorrected, Kruskal-Wallis test.

**Fig S4.**
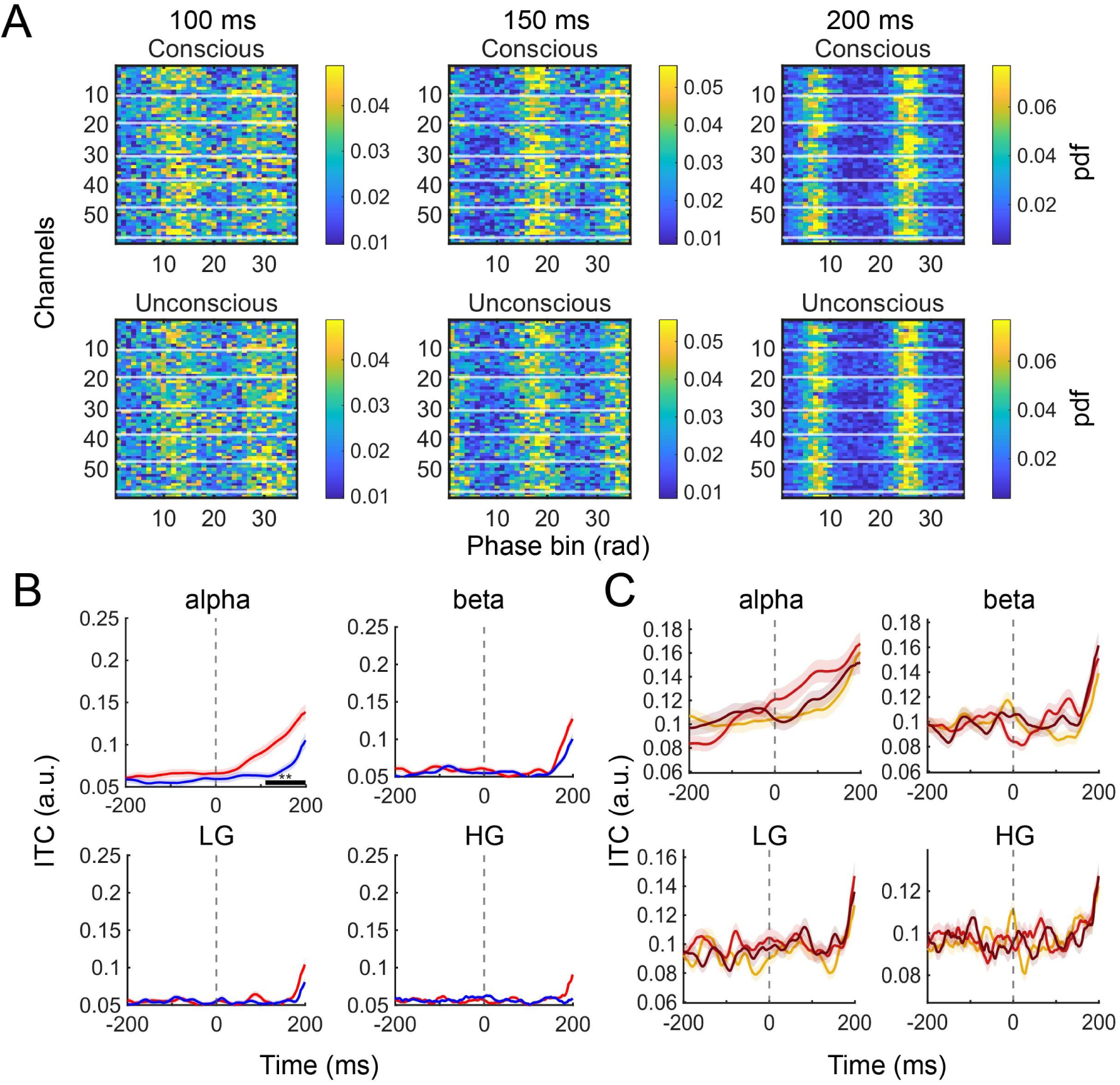
Phase alignment in cTha preceding conscious processing. (A) Heatmaps of phase distributions across individual recording sites at 0,100,200 ms. (B) ITC at alpha and the beta, low-gamma, high-gamma bands in cTha. (C) Different band-limited ITC across stimulus contrast groups in cTha during conscious trials.

**Fig. S5.**
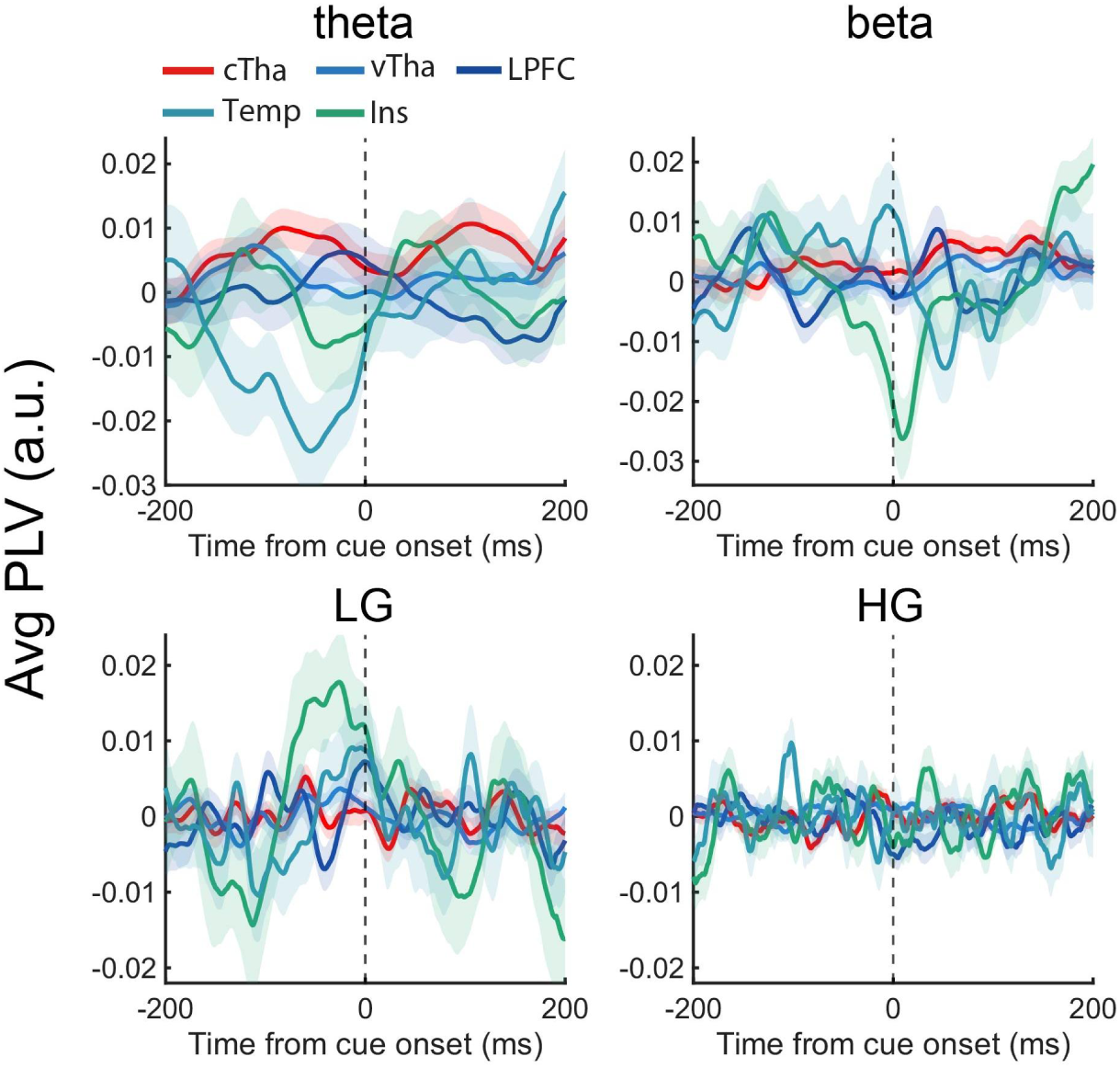
Early consciousness-related PLV changes in theta/alpha/low-gamma/high-gamma bands.

**Fig S6.**
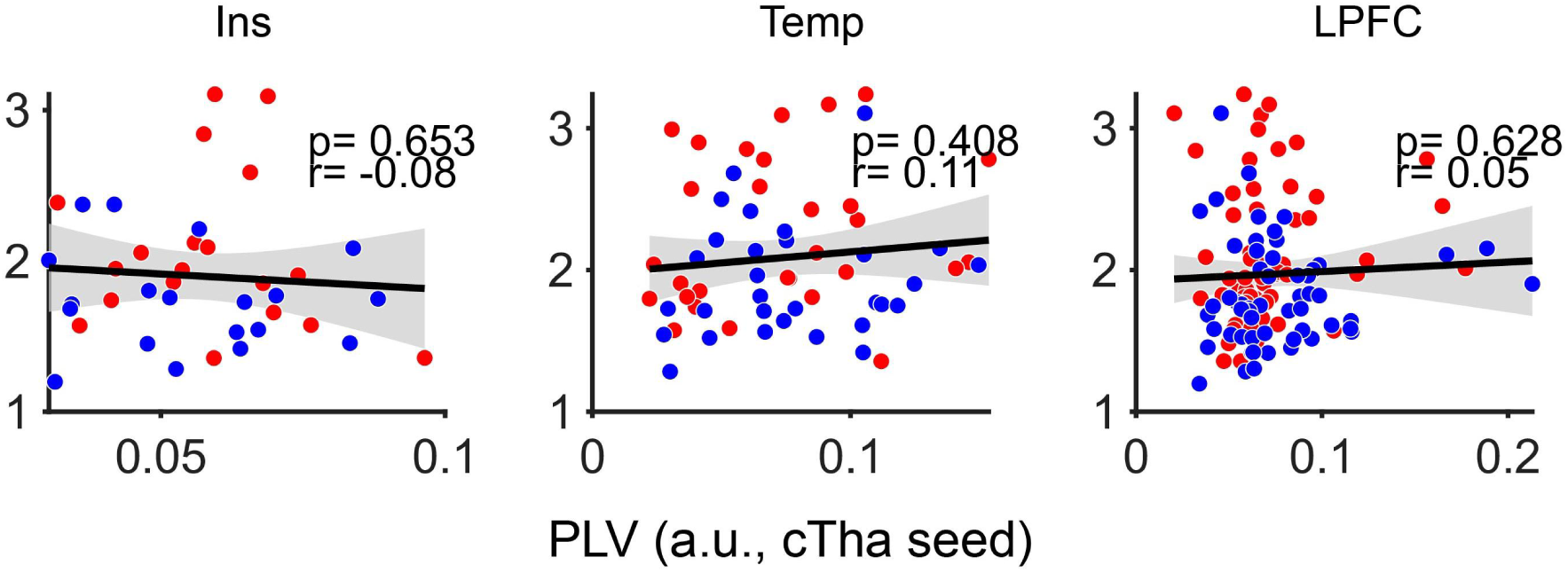
Correlations between early PLV and subsequent response magnitude (200-650 ms) for cTha-Ins (left), cTha–Temp (middle) and cTha–LPFC (right) coupling.

## Notes

### Competing Interest Statement

The authors have declared no competing interest.

## Reference

1. M. Nentwich, M. Leszczynski, C. E. Schroeder, S. Bickel, L. C. Parra, Intrinsic dynamic shapes responses to external stimulation in the human brain. eLife 14, RP104996 (2025).

2. E. Balestrieri, N. A. Busch, in J Neurosci. (United States, 2022), vol. 42, pp. 5058–5069.

3. J. Samaha, L. Iemi, S. Haegens, N. A. Busch, Spontaneous Brain Oscillations and Perceptual Decision-Making. Trends Cogn Sci 24, 639–653 (2020).

4. S. Dehaene, J. P. Changeux, Experimental and theoretical approaches to conscious processing. Neuron 70, 200–227 (2011).

5. F. Crick, C. Koch, Consciousness and neuroscience. Cereb Cortex 8, 97–107 (1998).

6. G. Northoff, F. Zilio, J. Zhang, Beyond task response-Pre-stimulus activity modulates contents of consciousness. Phys Life Rev 49, 19–37 (2024).

7. F. Di Gregorio et al., Tuning alpha rhythms to shape conscious visual perception. Curr Biol 32, 988–998.e986 (2022).

8. E. Balestrieri, N. A. Busch, Spontaneous Alpha-Band Oscillations Bias Subjective Contrast Perception. The Journal of Neuroscience 42, 5058–5069 (2022).

9. N. A. Busch, J. Dubois, R. VanRullen, The Phase of Ongoing EEG Oscillations Predicts Visual Perception. J Neurosci 29, 7869–7876 (2009).

10. L. Iemi et al., Ongoing neural oscillations influence behavior and sensory representations by suppressing neuronal excitability. NeuroImage 247, 118746 (2022).

11. C. Dembski, C. Koch, M. Pitts, Perceptual awareness negativity: a physiological correlate of sensory consciousness. Trends in Cognitive Sciences 25, 660–670 (2021).

12. G. Northoff, V. Lamme, Neural signs and mechanisms of consciousness: Is there a potential convergence of theories of consciousness in sight? Neurosci Biobehav Rev 118, 568–587 (2020).

13. G. A. Mashour, P. Roelfsema, J. P. Changeux, S. Dehaene, Conscious Processing and the Global Neuronal Workspace Hypothesis. Neuron 105, 776–798 (2020).

14. J. Forster, M. Koivisto, A. Revonsuo, ERP and MEG correlates of visual consciousness: The second decade. Conscious Cogn 80, 102917 (2020).

15. J. M. Shine, L. D. Lewis, D. D. Garrett, K. Hwang, The impact of the human thalamus on brain-wide information processing. Nat Rev Neurosci 24, 416–430 (2023).

16. Z. Fang et al., Human high-order thalamic nuclei gate conscious perception through the thalamofrontal loop. Science 388, eadr3675 (2025).

17. B. J. He, Spontaneous and Task-Evoked Brain Activity Negatively Interact. The Journal of Neuroscience 33, 4672–4682 (2013).

18. C. Babiloni, F. Vecchio, A. Bultrini, G. Luca Romani, P. M. Rossini, Pre- and Poststimulus Alpha Rhythms Are Related to Conscious Visual Perception: A High-Resolution EEG Study. Cerebral Cortex 16, 1690–1700 (2005).

19. J. Liang, Z. Yang, C. Zhou, Excitation–Inhibition Balance, Neural Criticality, and Activities in Neuronal Circuits. The Neuroscientist 31, 31–46 (2025).

20. K. B. Hengen, W. L. Shew, Is criticality a unified setpoint of brain function? Neuron 113, 2582–2598.e2582 (2025).

21. D. Toker et al., Consciousness is supported by near-critical slow cortical electrodynamics. Proceedings of the National Academy of Sciences 119, e2024455119 (2022).

22. D. Toker et al., Criticality supports cross-frequency cortical-thalamic information transfer during conscious states. eLife 13, e86547 (2024).

23. K. B. Hengen, W. L. Shew, Is criticality a unified setpoint of brain function? Neuron, (2025).

24. Z. Fang et al., The involvement of the human prefrontal cortex in the emergence of visual awareness. eLife 12, RP89076 (2024).

25. Z. Fang et al., Intracranial neural representation of phenomenal and access consciousness in the human brain. NeuroImage 297, 120699 (2024).

26. F. Stockart et al., Neural correlates of perceptual consciousness from within: A narrative review of human intracranial research. eLife 15, RP109604 (2026).

27. A. K. Seth, T. Bayne, Theories of consciousness. Nat Rev Neurosci 23, 439–452 (2022).

28. S. I. Kronemer et al., Human visual consciousness involves large scale cortical and subcortical networks independent of task report and eye movement activity. Nat Commun 13, (2022).

29. J. K. Hesse, D. Y. Tsao, A new no-report paradigm reveals that face cells encode both consciously perceived and suppressed stimuli. Elife 9, (2020).

30. S. Liu, Q. Yu, P. U. Tse, P. Cavanagh, Neural Correlates of the Conscious Perception of Visual Location Lie Outside Visual Cortex. Curr Biol 29, 4036–4044 e4034 (2019).

31. L. Fisch et al., Neural “ignition”: enhanced activation linked to perceptual awareness in human ventral stream visual cortex. Neuron 64, 562–574 (2009).

32. T. P. Reber et al., Single-Neuron Correlates of Conscious Perception in the Human Medial Temporal Lobe. Curr Biol 27, 2991–+ (2017).

33. J. Wilting, V. Priesemann, Inferring collective dynamical states from widely unobserved systems. Nat Commun 9, 2325 (2018).

34. V. A. F. Lamme, Challenges for theories of consciousness: seeing or knowing, the missing ingredient and how to deal with panpsychism. Philos Trans R Soc Lond B Biol Sci 373, (2018).

35. G. Tononi, M. Boly, M. Massimini, C. Koch, Integrated information theory: from consciousness to its physical substrate. Nat Rev Neurosci 17, 450–461 (2016).

36. B. R. Munn et al., A thalamocortical substrate for integrated information via critical synchronous bursting. P Natl Acad Sci USA 120, (2023).

37. J. O’Byrne, K. Jerbi, How critical is brain criticality? Trends in Neurosciences 45, 820–837 (2022).

38. A. I. Luppi et al., in Nat Neurosci. (United States, 2026).

39. A. S. Greene, C. Horien, D. Barson, D. Scheinost, R. T. Constable, Why is everyone talking about brain state? Trends in Neurosciences 46, 508–524 (2023).

40. E. Podvalny, L. E. King, B. J. He, Spectral signature and behavioral consequence of spontaneous shifts of pupil-linked arousal in human. Elife 10, (2021).

41. L. Waschke, S. Tune, J. Obleser, Local cortical desynchronization and pupil-linked arousal differentially shape brain states for optimal sensory performance. Elife 8, (2019).

42. R. Xia, X. Chen, T. A. Engel, T. Moore, Common and distinct neural mechanisms of attention. Trends in cognitive sciences, (2024).

43. P. Fries, Rhythmic attentional scanning. Neuron 111, 954–970 (2023).

44. J. Ashburner, K. Friston, Spatial transformation of images. spm course notes, (1997).

45. B. Fischl, FreeSurfer. Neuroimage 62, 774–781 (2012).

46. J. E. Iglesias et al., A probabilistic atlas of the human thalamic nuclei combining ex vivo MRI and histology. Neuroimage 183, 314–326 (2018).

47. A. Delorme, S. Makeig, EEGLAB: an open source toolbox for analysis of single-trial EEG dynamics. Journal of Neuroscience Methods 134, 9–21 (2004).

48. G. T. Einevoll, C. Kayser, N. K. Logothetis, S. Panzeri, Modelling and analysis of local field potentials for studying the function of cortical circuits. Nat Rev Neurosci 14, 770–785 (2013).

49. F. Tadel, S. Baillet, J. C. Mosher, D. Pantazis, R. M. Leahy, Brainstorm: a user-friendly application for MEG/EEG analysis. Comput Intell Neurosci 2011, 879716 (2011).

50. F. P. Spitzner et al., MR. Estimator, a toolbox to determine intrinsic timescales from subsampled spiking activity. PLOS ONE 16, e0249447 (2021).

